# Chromatin activity at GWAS loci identifies T cell states driving complex immune diseases

**DOI:** 10.1101/566810

**Authors:** Blagoje Soskic, Eddie Cano-Gamez, Deborah J. Smyth, Wendy C. Rowan, Nikolina Nakic, Jorge Esparza-Gordillo, Lara Bossini-Castillo, David F. Tough, Christopher G. C. Larminie, Paola G. Bronson, David Wille, Gosia Trynka

## Abstract

Complex immune disease variants are enriched in active chromatin regions of T cells and macrophages. However, whether these variants function in specific cell states or stages of cell activation is unknown. We stimulated T cells and macrophages in the presence of thirteen different cytokine cocktails linked to immune diseases and profiled active enhancers and promoters together with regions of open chromatin. We observed that T cell activation induced major chromatin remodelling, while additional exposure to cytokines fine-tuned the magnitude of these changes. Therefore, we developed a new statistical method that accounts for subtle changes in chromatin landscape to identify SNP enrichment across cell states. Our results point towards the role of immune disease variants in early rather than late activation of memory CD4+ T cells, and with limited differences across polarizing cytokines. Furthermore, we demonstrate that inflammatory bowel disease variants are enriched in chromatin regions active in Th1 cells, while asthma variants overlap regions active in Th2 cells. We also show that Alzheimer’s disease variants are enriched in different macrophage cell states. Our results represent the first in-depth analysis of immune disease variants across a comprehensive panel of activation states of T cells and macrophages.

## Introduction

Functional interpretation of complex disease variants is challenging as the majority of loci mapped through genome wide association studies (GWAS) reside in non-coding regions of the genome. Multiple studies have mapped GWAS variants to regulatory elements such as open chromatin regions and regions tagged by histone modifications^1–5^, implicating their role in gene expression regulation. The functional impact of non-coding GWAS variants is notoriously difficult to deconvolute, and may be specific to a particular cell type as well as a cell state context, such as different stages of cell activation^6^. Integrating GWAS variants with cell type specific chromatin marks can provide insights into disease causal cell types^1,4,7^. This approach has previously identified CD4+ T cells^4,8^ and monocytes^6,9^ as relevant cell types in the pathobiology of various complex immune diseases.

CD4+ T cells are key regulators of immune response and are crucial in the protection against pathogens. One of the hallmarks of CD4+ T cells is their plasticity; in particular, the ability to differentiate into a range of cell states in response to environmental signals. CD4+ T cells undergo initial activation when they recognise antigen displayed by antigen-presenting cells (APCs) in the context of co-stimulatory signals. Subsequently, activated T cells undergo proliferation and can be driven to differentiate into distinct T helper (Th) phenotypes, depending on the specific cytokines secreted by APCs. The major Th types include Th1, Th2, Th17 and induced regulatory T cells (iTregs), each exerting different functions in the immune response. Effector Th phenotypes are defined by the specific cytokines that they secrete, which in turn instruct other immune cells to acquire different phenotypes. For example, the Th1 cytokine IFN-γ polarizes macrophages to a proinflammatory (M1) phenotype with increased pathogen killing ability, while the Th2 cytokine IL-4 induces a tissue remodelling macrophage phenotype (M2)^10^. As such, the proper differentiation of T cells and macrophages following cytokine signals is a crucial step in eliciting an appropriate immune response.

Although it is established that immune disease variants localize to chromatin regions specific to CD4+ T cells and monocytes, it is not yet known if immune disease variants are further enriched in chromatin regions specific for a particular cytokine-induced cell state. To identify whether immune disease variants regulate cellular responses to cytokine polarization, we profiled chromatin accessibility using ATAC-seq, and active enhancers and promoters marked by H3K27ac (see **Methods**) in naive and memory CD4+ T cells as well as macrophages across 55 cell activation states, including early and late responses to activation and cytokine polarization (**Table S1**). We developed a new statistical method for assessing SNP enrichment in chromatin marks to point towards the effects of immune disease variants in specific cell states.

## Results

### Overview of the experimental design

The GWAS link to CD4+ T cells places this cell type at the heart of dysregulated immune responses in disease pathobiology. Key steps in regulating the quality of an immune response include the initial activation and differentiation of CD4+ T cells and the subsequent interaction of polarized T cells with downstream effector cells such as macrophages, whose activity is regulated by T cell-derived factors. In this study we focused on dissecting the role of immune diseases variants in regulating this circuitry. For this purpose, we stimulated monocyte-derived macrophages with T-cell-produced cytokines associated with inflammation and autoimmunity, including IFNγ, TNFα, IL-4, IL-23 and IL-26 (**Table S1**). Since macrophages are part of the fast-responding innate immune system, we measured cytokine induced activation at six hours (early response) and 24 hours (late response) and profiled the chromatin regulatory landscape. To mimic T cell activation *in vitro*, we stimulated T cells by delivering T cell receptor (TCR) and CD28 signals using beads coated with anti-CD3 and anti-CD28 antibodies. In addition, we exposed cells to cytokine cocktails promoting differentiation towards Th1, Th2, Th17 or iTreg cell fates, or to individual cytokines relevant to autoimmunity (IL-10, IL-21, IL-27, TNFα, and IFNβ)^11–15^ (**Table S1** and **Methods**). These stimuli were applied to memory and naive CD4+ T cells, which constitute the two major subsets of CD4+ T cells. We treated naive and memory cells separately because the two subsets differ in their response to stimulation^16^. Given that the response to stimulation develops over time^17^, we profiled T cells during both early and late activation. We defined early response as 16 hours in order to capture gene expression regulation prior to the first cell division. For late response we chose five days, which is when T cells acquire a defined effector phenotype. At each time point we profiled the chromatin regulatory landscape by assaying chromatin activity (H3K27ac ChIPmentation-seq) and chromatin accessibility (ATAC-seq). We then integrated these chromatin profiles with disease-associated variants to identify the most disease-relevant cell states.

### Accounting for peak properties refines cell-type specific disease SNP enrichment

We first used H3K27ac and ATAC-seq reads to identify active chromatin elements (peaks). We observed that stimulation induced thousands of new peaks across all cell types (**Figure 1B**). We then asked whether these peaks were the same across multiple cell states. To do this, we merged all peaks in naive and memory cell states across all time points into a common set of peaks. We repeated the same procedure for the macrophage data. We observed that the majority of peaks were shared between cell activation states. For example, in T cells, only 2% of H3K27ac and 0.8% of ATAC-seq peaks were condition specific (**Figure 1C**). We then quantified the level of activity of each element by using the number of reads in each peak. We found that quantitative levels of chromatin activity were highly variable across different cytokine induced cell states, as shown by the coefficient of variation of reads within peaks (**Figure 1C**). Importantly, the levels of chromatin accessibility were substantially less variable compared to H3K27ac which is in line with other reports showing that chromatin activity marked by H3K27ac is more informative in discriminating between closely related cell states than chromatin accessibility^18,19^ (**Figure 1C**).

**Figure 1.**
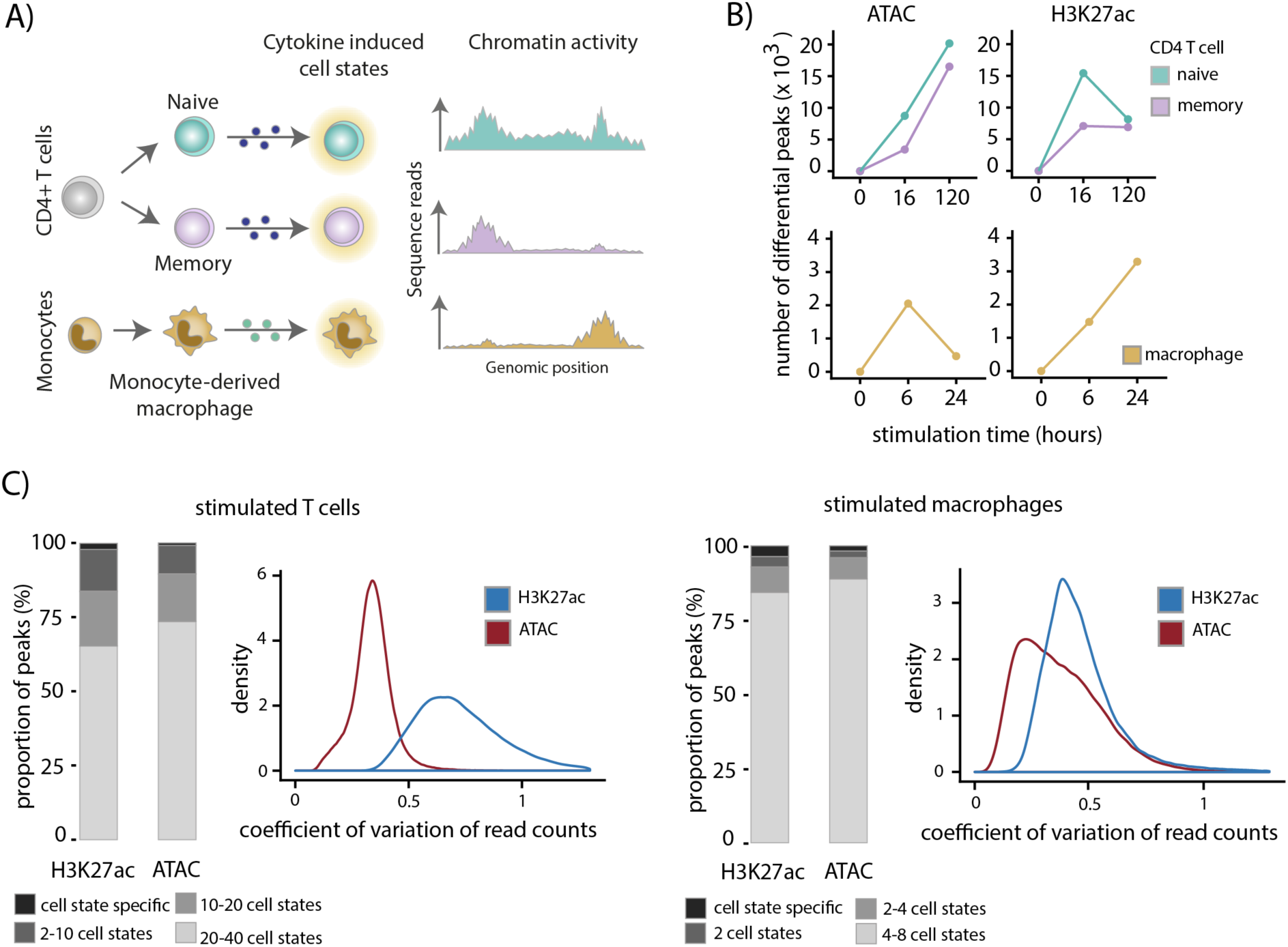
Quantitative changes in chromatin activity distinguish cytokine induced cell states. **A)** Overview of the study design. Naive and memory CD4+ T cells were isolated from peripheral blood from three healthy blood donors and stimulated with anti-CD3/anti-CD28 T cell activation beads. Macrophages were derived from monocytes using M-CSF. All cell types were cultured in the presence of immune disease-relevant cytokines and their chromatin activity was profiled at early and late time point. **B)** Number of differential H3K27ac and ATAC peaks upon T cell activation with anti-CD3/anti-CD28 T cell activation beads and macrophage activation with TNFa. **C)** Proportion of activation induced peaks that are shared between all macrophage or T cell states. Different shades of gray represent the extent of sharing. Density plots show coefficient of variation of the number of reads within ATAC and H3K27ac peaks.

We next assessed if immune disease variants were enriched in any specific cell state in our data set. Typically, disease SNP enrichment analyses rely on the presence or absence of overlap between associated variants and regulatory regions^1,7,20^. In our data set the majority of peaks were shared across cell states and therefore such a binary SNP-peak overlap approach would be unsuitable to discriminate between the different cellular conditions (**Figure S1**). A similar observation was previously made using partitioning heritability analysis of neuropsychiatric and metabolic disorders in highly correlated chromatin annotations of different brain regions^5^. To assess the immune disease enrichment across the different T cell and macrophage states we developed a new SNP enrichment method, *CHEERS* (Chromatin Element Enrichment Ranking by Specificity). In addition to SNP-peak overlap, our method takes into account peak properties as reflected by quantitative changes in read counts within peaks, corresponding to variable levels of H3K27ac or chromatin accessibility (**Figure 2** and **Methods**). Briefly, we first construct a matrix of quantile normalized read counts across peaks detected in all cell states (**Figure 2A**). For each peak we generate specificity scores where a high score is assigned to a peak with a higher read count in that cell state compared to all other states. Within each cell state, peaks are then ranked based on their specificity scores (specificity rank; **Figure 2B**). To assess disease SNP enrichment across the different cell states, for each locus we use the index variant and identify variants in strong LD (R^2^>0.8). Next, we identify peaks that overlap with the associated variants (**Figure 2C**). Importantly, our method is peak-centric, i.e. a peak that overlaps multiple disease associated variants in a locus contributes to the final cell-type specificity score only once. Alternatively, if the associated variants within a locus intersect multiple peaks, each independent peak contributes to the final enrichment score. We then calculate the mean specificity rank of all peaks that intersect disease associated variants and infer the significance of the observed SNP enrichment in the cell type by comparing it to a theoretical distribution.

**Figure 2.**
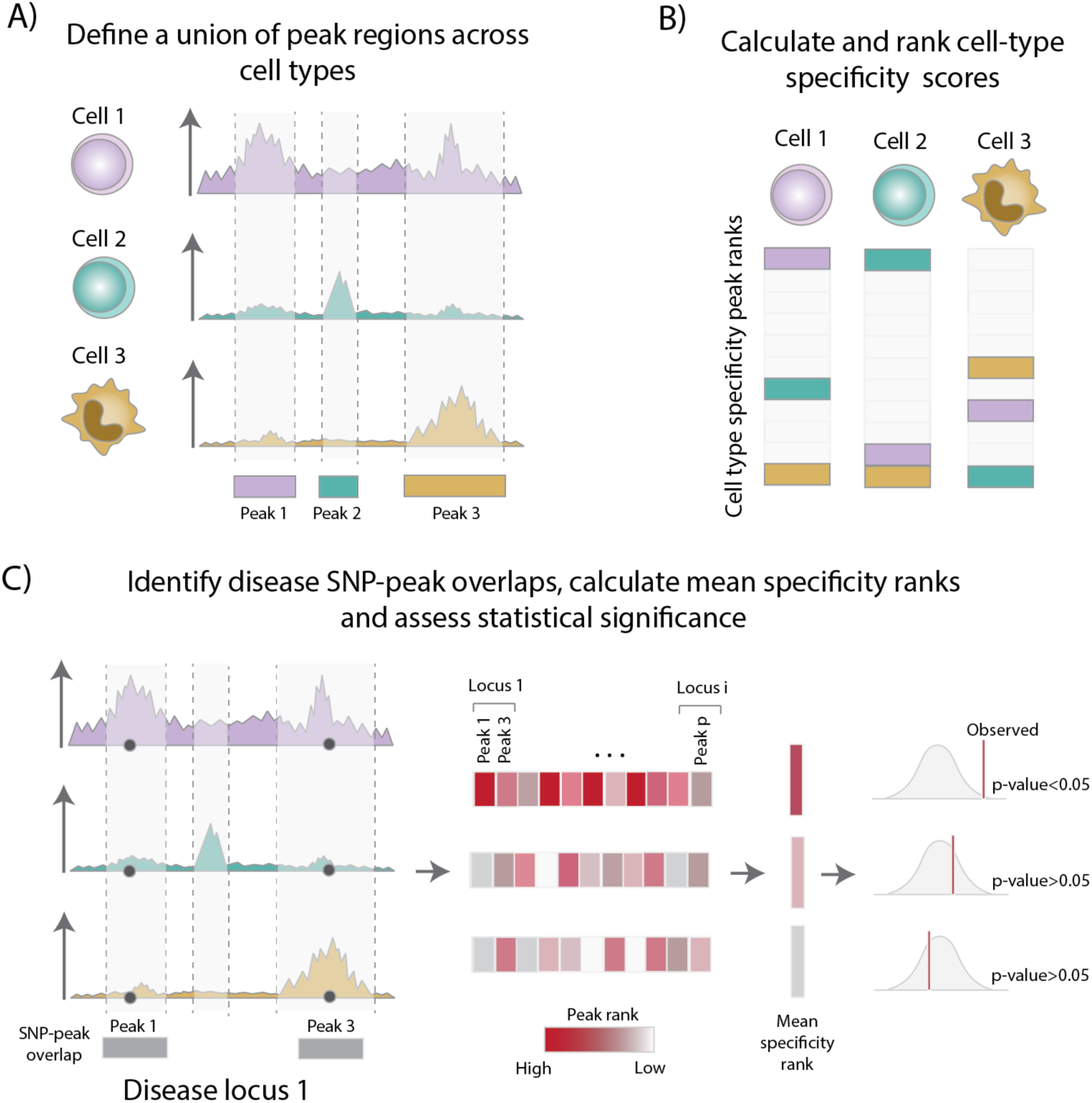
Overview of the *CHEERS* method. **A)** We first define a union of peak regions present across cell types and construct a matrix of normalized read counts. **B)** We then calculate and rank specificity scores. **C**) To test for disease SNP enrichment, we take all the index variants and identify variants in strong LD (r^2^ > 0.8). We then identify peaks that overlap with the associated variants, calculate the mean specificity ranks and assess statistical significance.

We benchmarked *CHEERS* by testing for immune disease SNP enrichment across cell types assayed with H3K27ac ChIP-seq as a part of the BLUEPRINT project^3^. Our method identified enrichment of multiple sclerosis (MS) variants in T and B cells, while inflammatory bowel disease (IBD) and rheumatoid arthritis (RA) variants showed enrichment exclusively in T cells (**Figure S2**). These results agree with previous reports^7,21,22^ and validate our method.

Next, we used simulations to assess the sensitivity of *CHEERS* (**Figure S3**). With over 23% of variants overlapping the top 10% most cell-type-specific peaks we observed over 80% power to detect a significant enrichment (p-value < 0.01). However, at least 78% SNPs needed to map within peaks ranked as low as 60%-70% of cell-type-specific peaks to achieve 80% power of detecting a significant enrichment. This indicates that *CHEERS* can detect a significant cell type enrichment when a sufficient proportion of trait associated variants overlap peaks with high cell-type-specificity ranks. Based on the results from the BLUEPRINT data and the simulations we concluded that *CHEERS* is able to identify enrichment in active chromatin elements across both diverse and closely related cell types.

### Immune disease SNPs are enriched in active chromatin elements specific to early activation of memory T cells

We applied *CHEERS* to our data from cytokine induced cell states to test for SNP enrichment across six diseases with an immune component, including multiple sclerosis (MS), rheumatoid arthritis (RA), inflammatory bowel disease (IBD), asthma, allergies and Alzheimer’s disease (AD) (**Table S2**). The tested traits included between 17 and 132 associated loci, resulting in between 26 (asthma) and 386 (IBD) SNP-peak overlaps contributing to the final enrichment (**Table S3 and Table S4**).

Across all tested traits, only AD variants were significantly enriched in macrophages (**Figure 3**). This agrees with recent reports implicating microglia, a subtype of macrophages present in the brain, in the pathogenesis of AD^23,24^. We observed that AD variants were enriched in active chromatin regions present in all macrophage conditions, and in both early and late activation. On the other hand, we found that risk loci for MS, RA, IBD, allergy and asthma were predominantly enriched in T cells.

**Figure 3.**
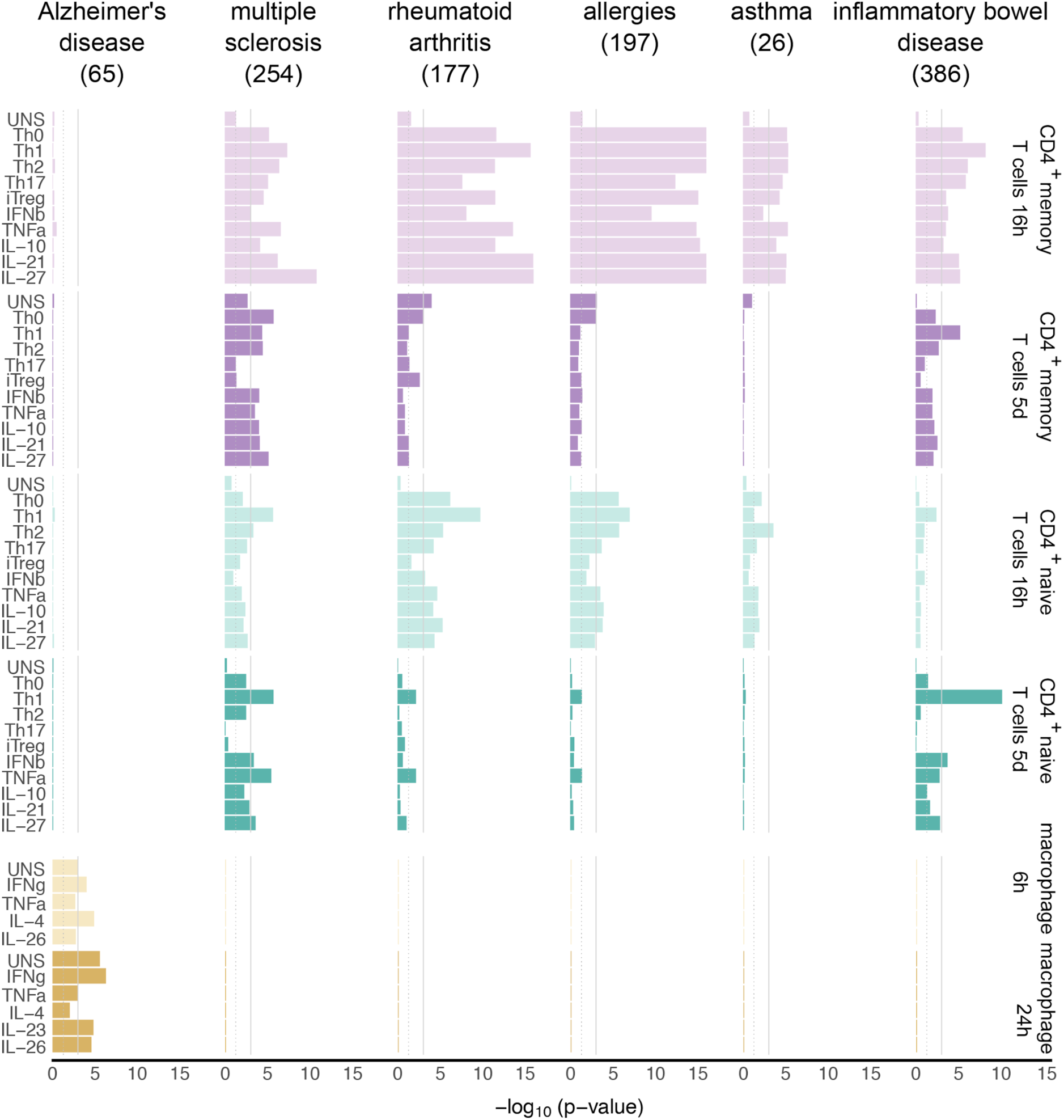
Disease SNP enrichment in H3K27ac regions in cytokine induced cell states. We used *CHEERS* to test for SNP enrichment in H3K27ac regions across cytokine induced cell states. The numbers in parentheses represent the number of SNP-peak overlaps per disease. The height of the bars represents -log10(p-value). The dotted gray line represents the nominal p-value threshold of 0.05, while the solid gray line represents the Bonferroni corrected p-value threshold of 0.00091 (corrected for the number of cell states in the study).

Surprisingly, we found that enrichment of immune disease loci was particularly strong in chromatin regions upregulated during early activation of memory T cells (16 hours; **Figure 3**). In contrast, resting memory T cells, which were cultured without any stimulus, showed no significant enrichment in any of the tested diseases, suggesting that it is specifically the regulation of cell activation which drives this enrichment. To corroborate this, we explored the individual loci driving the enrichment and found a clear signal in proximity of genes involved in T cell activation. For example, RA GWAS variants overlapped a peak in the C-C Motif Chemokine Receptor 6 (*CCR6*) gene which showed higher levels of acetylation during early activation of memory and not naive T cells (**Figure 4A**). Another example included three RA associated variants that overlapped a peak in acyl-CoA Oxidase Like protein (*ACOXL*) that was highly upregulated only during early activation of both naive and memory T cells (**Figure 4A**). By using genes in proximity to variants overlapping the top 25% specific peaks driving the enrichment in early activation of memory T cells we observed enrichment of pathways such as T cell activation and differentiation, and leukocyte activation (**Figure 4B**). This suggests that immune disease variants overlap enhancers and promoters that regulate gene expression programmes underlying early activation of memory T cells. Furthermore, for MS, RA and IBD we also observed that the enrichment in early T cell activation varied between individual cytokine induced cell states. For example, across the cell states significantly enriched for MS variants, IL-27 stimulation showed the most significant enrichment (p=6.5×10^−12^). This is in concordance with previous studies that reported elevated levels of IL-27 in the cerebrospinal fluid of MS patients^25,26^.

**Figure 4.**
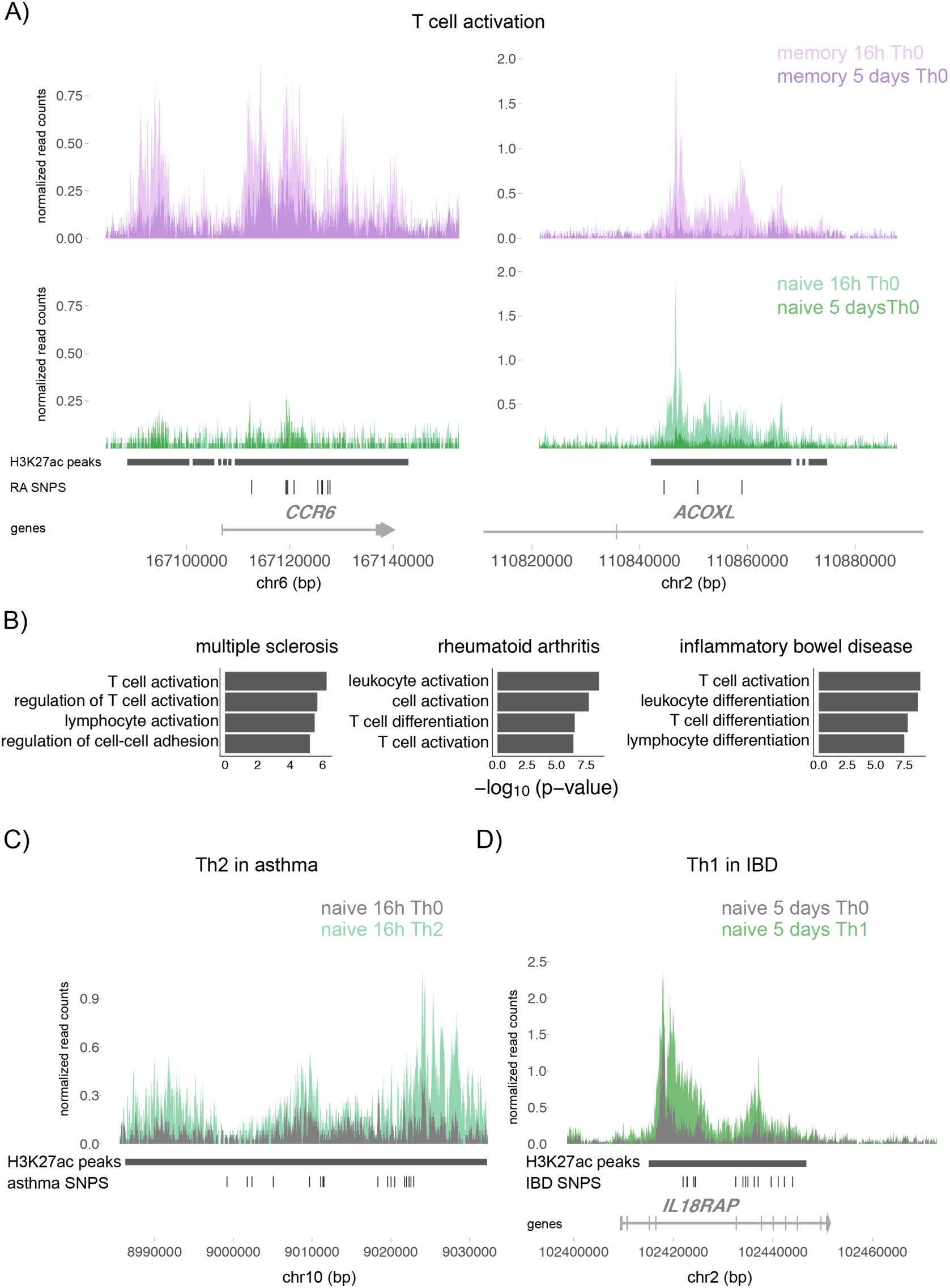
Example loci driving immune disease SNP enrichment in induced CD4+ T cell states. **A)** Examples of loci driving rheumatoid arthritis enrichment in early and not late activation of T cells. Shades of purple and green represent early and late memory and naive T cell activation respectively. **B)** Pathway enrichment analysis using all the genes in proximity of the 25% top specific peaks overlapping immune disease variants. Example loci driving the enrichment of **C)** asthma variants in naive CD4+ T cells 16h stimulation towards Th2 phenotype and **D)** IBD variants in naive CD4+ T cells after 5 days stimulation towards Th1 phenotype.

We also observed significant enrichment in early activation of naive CD4+ T cells; however the enrichment was less significant in comparison to memory cells. For example, in MS and RA, Th1 cells showed the strongest enrichment (p=2.08×10^−6^ and 1.84×10^−10^ respectively), while in asthma we observed significant enrichment only in early activation of naive CD4+ T cells polarized towards the Th2 phenotype (**Figure 3**), pointing towards the well-established role of Th2 cells in the pathobiology of asthma^27,28^. Among the loci driving the enrichment of asthma in Th2, we identified an active chromatin region located 900kb away from the hallmark Th2 gene, *GATA3*, and induced only upon Th2 polarization (**Figure 4C**).

Conversely, we observed a significantly lower signal in later stages of T cell activation (5 days). Here, the enrichment was apparent only in selected T cell states and diseases, for example, we observed a strong enrichment of IBD risk variants in naive and memory T cells polarized with Th1 cytokines (p=7×10^−11^ and p=5.91×10^−6^, respectively). Among the loci driving this signal was the *IL18RAP* locus, where a 30kb region undergoes substantial acetylation upon Th1 polarization (**Figure 4D**). In MS we also observed enrichment in late memory T cell activation, albeit to a lower extent than in early activation. On the other hand, we have not observed any enrichment in control traits such as bone mineral density, breast cancer and coronary artery diseases (**Figure S4**). Finally, we recapitulated similar results using our ATAC-seq data, although enrichment in ATAC-seq was weaker than in H3K27ac (**Figure S5)**. Globally, our results point towards a number of variants associated with immune mediated diseases that function in early activation of CD4+ T cells, predominantly in memory cells.

## Discussion

Identifying the most relevant cellular context in which disease associated variants function is critical for designing meaningful functional follow-up studies. Here, we used an *in vitro* system to identify cytokine induced cell states relevant in the pathobiology of immune-mediated diseases. We observed that most changes in the chromatin landscape of CD4+ T cells resulted from TCR and CD28 stimulation alone, while the presence of specific cytokines had a modulatory effect on the levels of these changes. As a consequence, T cell states had very similar chromatin landscapes and currently available SNP enrichment methods could not distinguish between them. To address this, we developed *CHEERS*, a statistical method that takes into account quantitative changes in chromatin activity.

Using *CHEERS*, we were able to refine the enrichment of immune disease variants observed in previous studies^7,21,22^ to specific cellular contexts. Across the five immune diseases that we tested, we observed that the associated variants were enriched in early activation of memory CD4+ T cells. The role of CD4+ effector memory T cells in conditions like RA has been previously suggested^21,29^. Additionally, a recent study which profiled open chromatin regions across subsets of immune cells in resting and stimulated states identified enrichment of immune disease variants in T cell activation^33^. Our study now provides further evidence for the importance of this cell type in the biology of complex immune mediated diseases and builds a case for dysregulation of specific cellular processes. Our results suggest that GWAS variants for immune-mediated diseases affect genetic regulation mostly during the initial phase of memory T cell activation, but less so during the effector response seen at the second time point. This emphasises the importance of tight regulation of T cell activation, suggesting that many subtle effects of immune variants lead to dysregulated early cell responses. This agrees with observations from severe immune disorders where, for instance, deficiency in the expression of one of the main regulators of T cell activation, CTLA-4, has been associated with the development of autoimmune diseases due to uncontrolled T cell activation^30–33^. Furthermore, it is worth noting that immune-mediated diseases are more often diagnosed in adults, which coincides with a shift in the frequency of T cells from predominantly naive to predominantly memory. Therefore our results suggest that focusing on regulation of early activation of memory CD4+ T cell may provide an important axis for development of new treatment options.

In addition to early activation of memory CD4+ T cells, we observed selected diseases where the associated variants were enriched in distinctive cytokine polarised cell states. For example, IBD risk variants were highly enriched in naive and memory T cells polarized with Th1 cytokines. This agrees with previous findings from immunophenotyping studies showing that lamina propria T cells express high levels of Stat4, T-bet and IL-12R, which are all induced by Th1 polarizing cytokines^34^. This could indicate that a proportion of IBD variants regulate the function of Th1 cells, suggesting the involvement of this cell type in IBD development. Another example was asthma, where we detected enrichment in naive T cells polarized towards the Th2 phenotype. Our results provide evidence of a genetic basis of the previous results from immunology studies, which implicate Th2 cells in allergic asthma^27,35^ and link disease associated variants to dysregulation of this cell type.

Across the tested immune diseases, we did not observe a significant enrichment in any of the macrophage stimulatory cell states, suggesting that autoimmune inflammation is mostly driven by T cells. In contrast, variants associated with AD point to a crucial role of macrophages in the disease. Recent studies have highlighted the role of microglia in AD^23,24,36^, a type of resident macrophage in the central nervous system (CNS). Our results suggest that a proportion of AD associated variants could be studied in an *in vitro* model of monocyte differentiated macrophage. This can have significant implications, as cells from the CNS are challenging to collect. Finally, the lack of stratification of AD variants between the different cytokine induced macrophage states could indicate that some AD variants might regulate more general functions of the macrophage lineage.

Our study is the first to systematically profile changes in chromatin regulatory landscape induced by cytokines during activation of human immune cells. This provides a valuable resource to identify appropriate cell models for studying how genetic variants lead to diseases.

## Authors contributions

GT and BS conceived and designed the project. BS, ECG and DJS carried out the experimental work. BS and ECG performed the data analysis. BS, ECG, GT, WCR, NN, JEG, DT, CL, DW, LBC and PGB interpreted the results. BS, DW, ECG and GT developed the *CHEERS* method. LBC calculated LD blocks. GT supervised the analysis. GT, BS, ECG, WR, NN, JEG, DT, CL, DW and PGB wrote the manuscript.

## Code and data availability

Upon acceptance we will make the data publicly available through the European Genome-Phenome Archive (EGA; http://ega-archive.org). CHEERS code is available through github (https://github.com/trynkaLab/CHEERS)

## Methods

### Sample processing and cell activation

Blood samples were obtained from three healthy individuals. The human biological samples were sourced ethically and their research use was in accord with the terms of the informed consents under an IRB/EC approved protocol (15/NW/0282). Peripheral blood mononuclear cells (PBMCs) were isolated using Ficoll-Paque PLUS (GE healthcare, Buckingham, UK) density gradient centrifugation. Naive and memory CD4+ T cells were isolated from PBMCs using EasySep® naive CD4+ T cell isolation kit and memory CD4+ T cell enrichment kit (StemCell Technologies, Meylan, France) according to the manufacturer’s instructions. T cells were stimulated with anti-CD3/anti-CD28 human T-Activator Dynabeads® (Invitrogen) at 1 bead : 2 T cell ratio. Cytokines were added at the same time as the stimulus and cells were harvested after 16h and 5 days (for the list of cytokines, and their concentration refer to **Table S1**).

Monocytes were isolated using EasySep® monocytes isolation kit according to the manufacturer’s instructions. In order to generate macrophages, monocytes were plated in a 100mm x 20mm dish and cells were treated with 800 U/ml M-CSF (PeproTech) for seven days. Following macrophage differentiation, they were stimulated with cytokines for 6 and 24 hours (for the list of cytokines, and their concentration refer to **Table S1**).

### ChIPmentation-seq

In order to profile active enhancers and promoters, cells were washed with RPMI and Dynabeads® were removed using a DynaMag® magnet (Thermo Fisher). Next, cells were resuspended at 1×10^6^ cells/ml in FACS buffer (PBS buffer supplemented with 10% FCS and 1 mM EDTA) and chromatin was cross-linked by adding 1% formaldehyde and incubating at 37°C for five minutes. To quench the reaction we added glycine and cells were washed in cold PBS buffer. Cross-linked cell pellets were frozen in liquid nitrogen and stored at −80°C until further processing.

To perform ChIPmentation, cross-linked pellets were thawed and processed using the iDeal® ChIP-seq kit for histones (Diagenode) according to the manufacturer’s instructions. Briefly, cells were lysed and sonicated, and the resulting material was used for immunoprecipitation (IP). Sonication was performed using a Bioruptor® Pico (Diagenode), with 6 sound pulse cycles of 30 seconds each. For immunoprecipitation (IP), we used ChIP grade antibodies specific to human H3K27ac histones (Diagenode). A negative control undergoing no IP (input) was also generated for each cell type. After IP, the recovered chromatin fragments were tagmented as previously described ^37^. Briefly, 2 μl of IP material were resuspended in 30 ul ChIP-seq buffer (Diagenode) supplemented with 1 μl Tn5 (Illumina). Samples were then reverse cross-linked using the iDeal® ChIP-seq kit for histones according to the manufacturer’s instructions. The resulting DNA was purified using SPRI magnetic beads (AMPure XP A63881 Beckman Coulter). Enrichment of active chromatin regions was verified by qPCR.

Sequencing libraries were constructed from the obtained material using the Nextera® DNA library preparation kit according to the manufacturer’s instructions. Briefly, DNA was amplified by PCR and fragments of inappropriate sizes were removed using Agencourt AMPure XP beads (BD). Finally, samples were pooled and loaded into an Illumina® HiSeq 2500 instrument for paired-end sequencing. In order to minimise batch effects, samples were randomized before sequencing. We obtained on average 63 million paired-end reads per sample.

### ATAC-seq

In order to profile open chromatin regions, stimulated cells were washed with RPMI and Dynabeads® were removed using a DynaMag® magnet (Thermo Fisher). Next, tagmentation was performed using the fast ATAC protocol ^38^. Briefly, 50,000 cells were washed in cold PBS buffer and resuspended in 50 μl of Nextera® tagmentation buffer supplemented with 0.01% digitonin and 2.5 μl Tn5 transposase (Nextera). Samples were then incubated at 37°C and 800 rpm for 30 minutes. After tagmentation, DNA was purified using MinElute® PCR columns (Qiagen) according to the manufacturer’s instructions and stored at −80°C until library preparation.

Sequencing libraries were generated from tagmented DNA using the Nextera® DNA library preparation kit according to the manufacturer’s instructions. Briefly, DNA was amplified by PCR and fragments of inappropriate sizes were removed using Agencourt AMPure XP beads (BD). Finally, samples were pooled and loaded into an Illumina® HiSeq 2500 for paired-end sequencing. In order to minimise batch effects, samples were randomized before library preparation and before sequencing. We obtained on average 58 million paired-end reads per sample.

### ATAC-seq and ChM-seq analysis

We assessed the quality of reads using fastx and trimmed the adapters using skewer (v0.2.2)^39^. We then mapped reads to the human genome reference GRCh38 using bwa mem (v0.7.9a)^40^ only kept uniquely mapped reads. We also removed PCR duplicates and mitochondrial reads in ATAC-seq using samtools (v0.1.9)^40,41^. This resulted in final BAM files containing uniquely mapped, non-mitochondrial reads that were used for peak calling. Finally, we calculated insert size distributions using PICARD tools (v2.6.0) to remove samples with over- or under-sonicated chromatin and removed samples with a skewed distribution of insert sizes.

Peaks were called using MACS2 (v2.1.1)^42^ -q 0.05 --nomodel --extsize 200 --shift −100 for ATAC; and --broad --broad-cutoff 0.1 --nomodel --extsize 146 for H3K27ac ChM.

Next, we calculated the fraction of reads in peaks (FRiP) to investigate the quality of our data. The average FRiP for H3K27ac ChM was 46,64% and for ATAC was 17.03%. Samples with FRIP < 5%, skewed distribution of insert sizes, or < 20 million QCed reads were removed from the downstream analysis.

To call peaks per cell state, QCed BAM files corresponding to biological replicates of the same condition were merged using samtools and peaks were called as described above but with two additional parameters. First, when pooling reads across individuals and within the cell state we expected to see a proportion of identical sequence reads in independent donors by chance, therefore we used the --keep-dup flag in MACS2, as PCR duplicated reads had already been removed. Second, we increased the q-value threshold to 0.1. In ATAC-seq, where the quality of the data was lower compared to ChM, we wanted to ensure that noisy peaks were excluded from our enrichment analysis. Therefore we used stringent parameters and kept all peaks with fold enrichment > 4 and q-value < 10^−4^. In ChM, where peaks are broad and FRiP values higher, we kept all peaks with fold enrichment > 2 and q-value < 10^−2^. This resulted in the final BED files which were used used for GWAS enrichment analysis. To define differentially accessible regions and differentially modified histone regions, we used DESeq2. We compared all conditions to the resting state or to Th0 and used Benjamini-Hochberg controlled FDR of 5% and an absolute fold-change ≥ 1.

#### Disease enrichment with GoShifter

We ran GoShifter on ATAC and H3K27ac peaks as described previously^7^. We ran 10,000 permutations. Command used: *python goshifter.py -s SNP_list -a annotation/file/path -p 10000 -l LD/file/path -o output_name*.

### Chromatin Element Enrichment Ranking Specificity (CHEERS)

We first merged the peaks across all cell types and cell states using bedtools (v2.22.0) merge option in order to get a unified set of peaks. Then, for each cell type and cell state we quantified the the number of reads within the merged peak regions using featureCounts (v1.5.1)^43^. We normalized each peak for the library size by scaling the peak read counts to the sample with the greatest count of informative sequence reads:

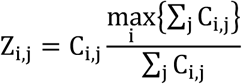

where *Ci, j* is the number of reads falling within peak *j* in cell state *i*

To ensure that our analysis was not confounded by the low confidence peaks, we removed the bottom 10th percentile of peaks with the lowest read counts and obtained a final count of 132,236 peaks for H3K27ac ChM and 740.221 for ATAC-seq.

In order to compare the peaks across the cell types and cell conditions, we also quantile normalized the library size-corrected peak counts. We then transformed the read count of each peak into a score that reflects cell type specificity (specificity score). For that, we divided the normalized read counts of each peak in each condition by the Euclidean norm for that peak across all conditions, as described in the following formula:

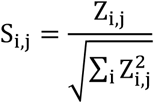

where *Si, j* is the specificity score, and *Zi, j* is the normalized number of reads within a peak *j* in a condition *i*. This score is a number from 0 to 1, where 1 means a peak has high read counts in only one cell state and 0 means the peak shows no read in that cell state.

Then, within each cell state, peaks were ranked by specificity score from the highest to the lowest score, where the peak least specific to the cell state was ranked 1. If multiple peaks have equal specificity scores, the same lower rank is assigned to all of them.

To test for disease SNP enrichment, we take all the index variants, identify variants in LD (r2>0.8) and overlap them with peaks. Importantly, our method is peak-centric, i.e. a peak that overlaps multiple disease associated variants in a locus contributes to the final cell-type specificity score only once. On the other hand, if within a locus the associated variants intersect multiple peaks, each independent peak contributes to the final enrichment score. We then calculate the mean specificity rank of all peaks which intersect disease associated variants. We inferred the significance of the observed enrichment by fitting a discrete uniform distribution. Within each cell state all ranks (1, 2, 3…*N*) can be observed with an equal probability and thus they follow a discrete uniform distribution (Kolmogorov-Smirnov test p=1) with mean (*μ*):

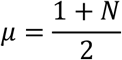

and variance:

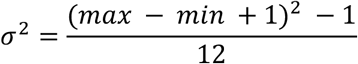

where *max* and *min* are the maximal and minimal ranks in the dataset. When we substitute *max* with number of peaks (*N*) and *min* with the minimal rank (1) we obtain the following formula:

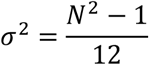

which, under the central limit theorem converges to:

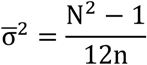

where 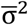 is the variance of the mean of the n peak ranks overlapping the GWAS SNPs assuming that these overlap at random.

Finally, we calculate p-values as:

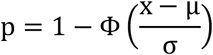

where the 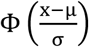 is a cumulative normal distribution for.x∼*N* (μ, σ^2^); x is the observed rank, and μ and σ are the expected values under the null hypothesis that all ranks occurred as random.

To ensure that this method accounts for any possible unidentified properties of the data, such as correlations, we also assessed the significance of the enrichment using an empirical, permutation based strategy. For that, within each of the tested cell states, we randomly sampled sets of peaks, matching for the number of peaks overlapping GWAS variants, and calculated the mean of their ranks. We repeated this process N times, assessed the frequency at which the mean of permuted ranks was greater or equal to the mean of observed ranks, and derived an empirical p-value:

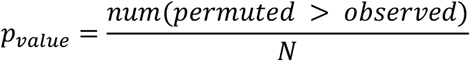

where N is a number of permutations. Both approaches yielded similar results while the p-values from CHEERS were not limited by the number of permutations (R^2^ = 0.96, p-value < 2.2e-16).

#### Power calculations for CHEERS

To estimate the power of *CHEERS*, we simulated 100 SNP-peak overlaps and tested for enrichment. Each simulation was repeated 100 times and power was estimated as the percentage of simulations that yielded a significant enrichment (p-value < 0.01). We designed the simulations such that a given percentage of SNPs (ranging from 0% to 100%), but not the remaining SNPs, always overlapped with peaks in the top 10th percentile of specificity. Finally, in order to test how specific the peaks needed to be for our method to detect enrichment, we repeated the simulations at lower specificity percentiles.

## Supplementary Figures

**Figure S1.**
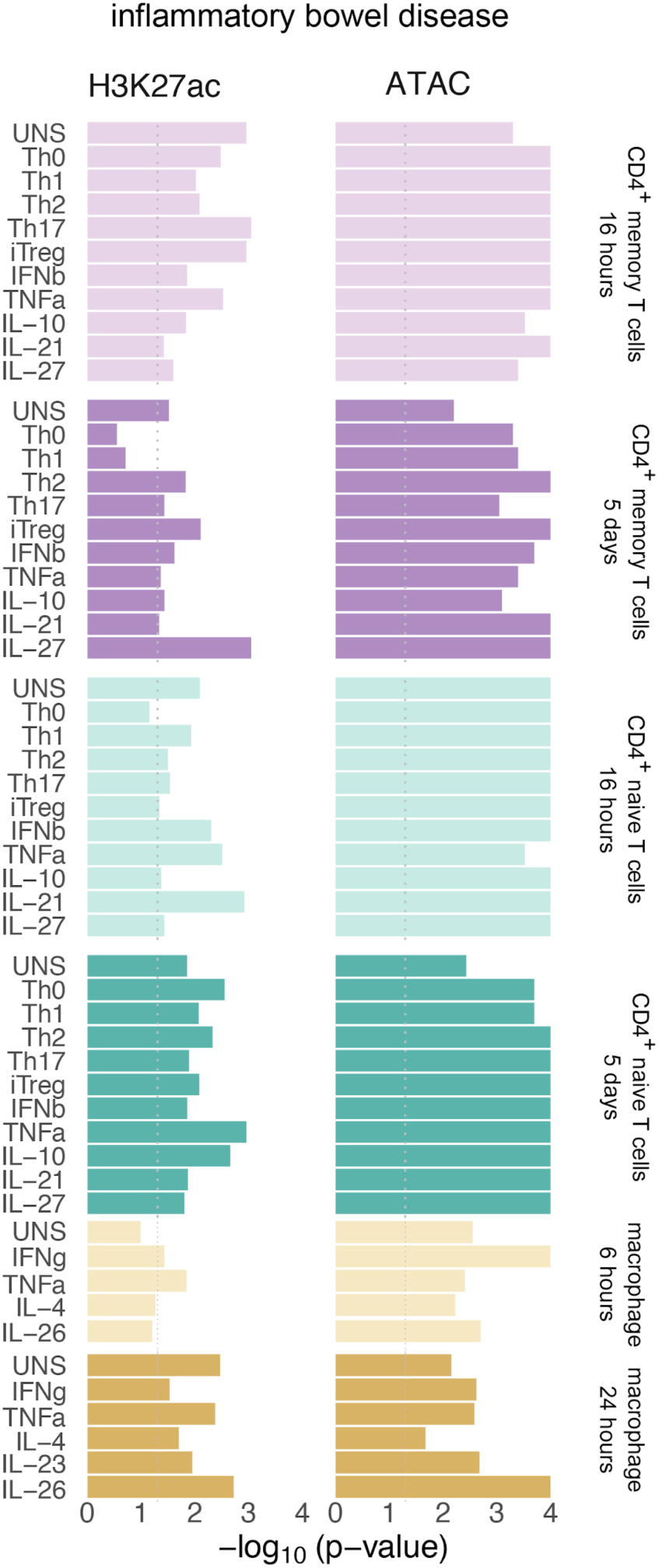
Binary SNP-peak overlap is not sufficient to discriminate between closely related cell states. We used GoShifter to test for SNP enrichment across cytokine induced cell states. We ran 10,000 permutations with default parameters.

**Figure S2.**
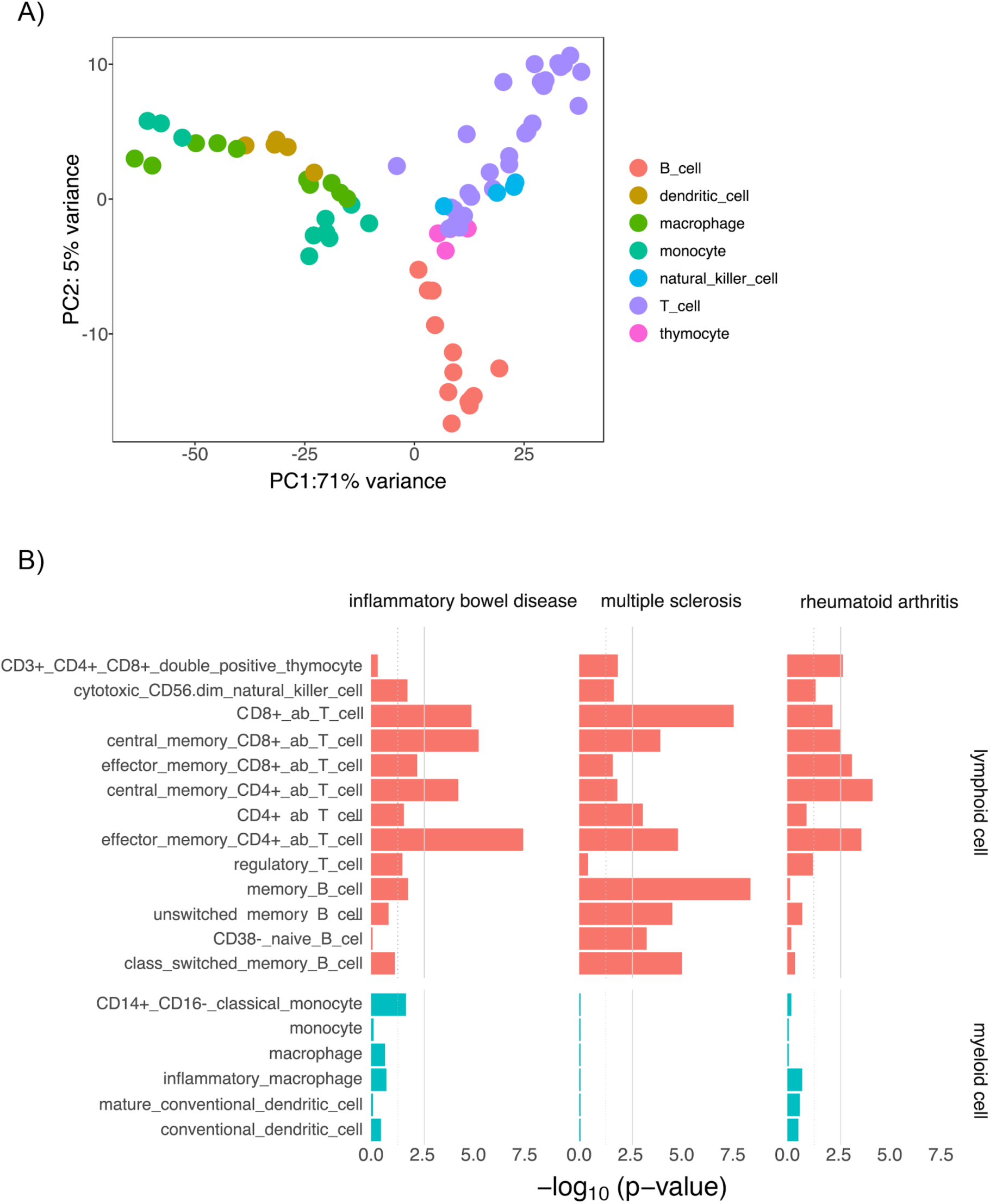
Disease SNP enrichment in H3K27ac regions in blood cell types. **A)** PCA of cell types assayed as a part of BLUEPRINT project. **B)** *CHEERS* results across different blood cell types from the BLUEPRINT project. The dotted gray line represents the nominal p-value threshold of 0.05, while the solid gray line represents the Bonferroni corrected p-value threshold of 0.0026 (corrected for the number of cell types tested).

**Figure S3.**
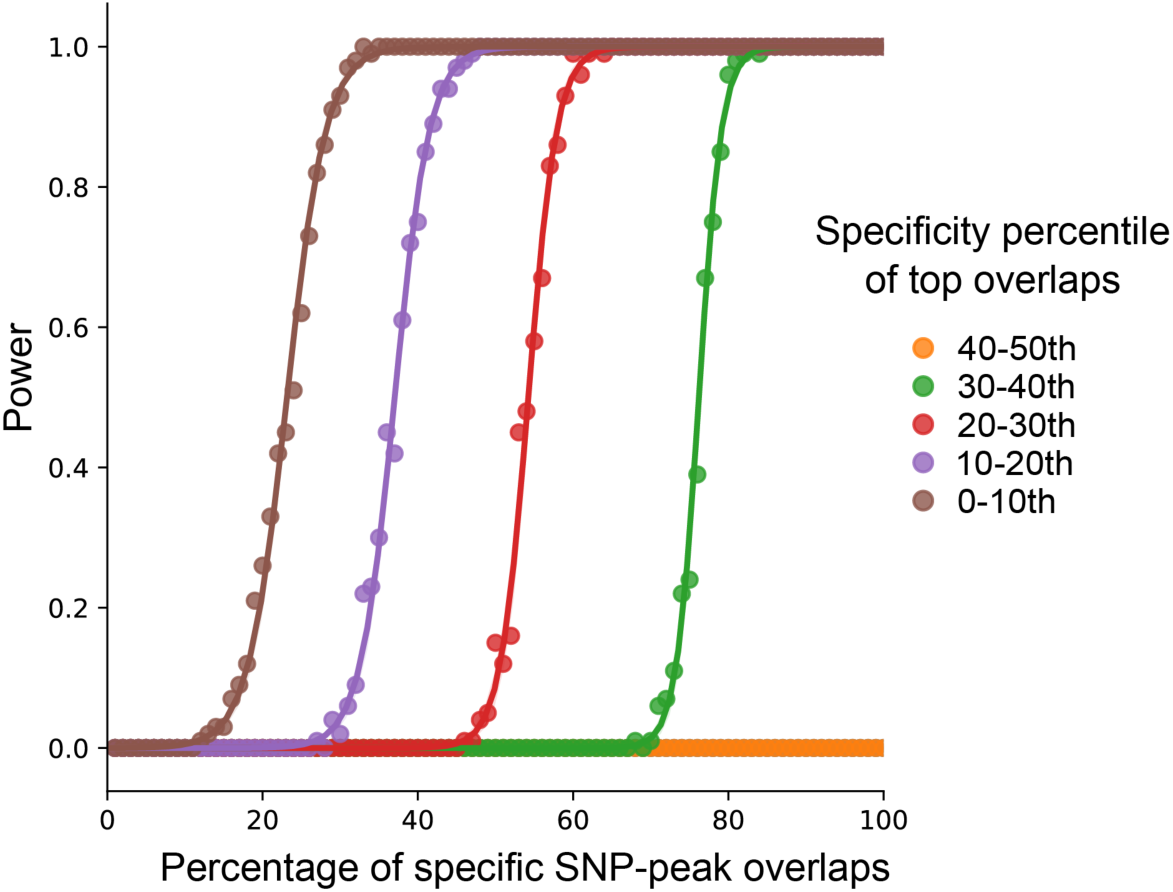
Power calculations for *CHEERS*. Graph shows the power to detect the enrichment (p< 0.01) with different percentage of specific SNP-peak overlaps. Different colours represent specificity percentile of top overlaps.

**Figure S4.**
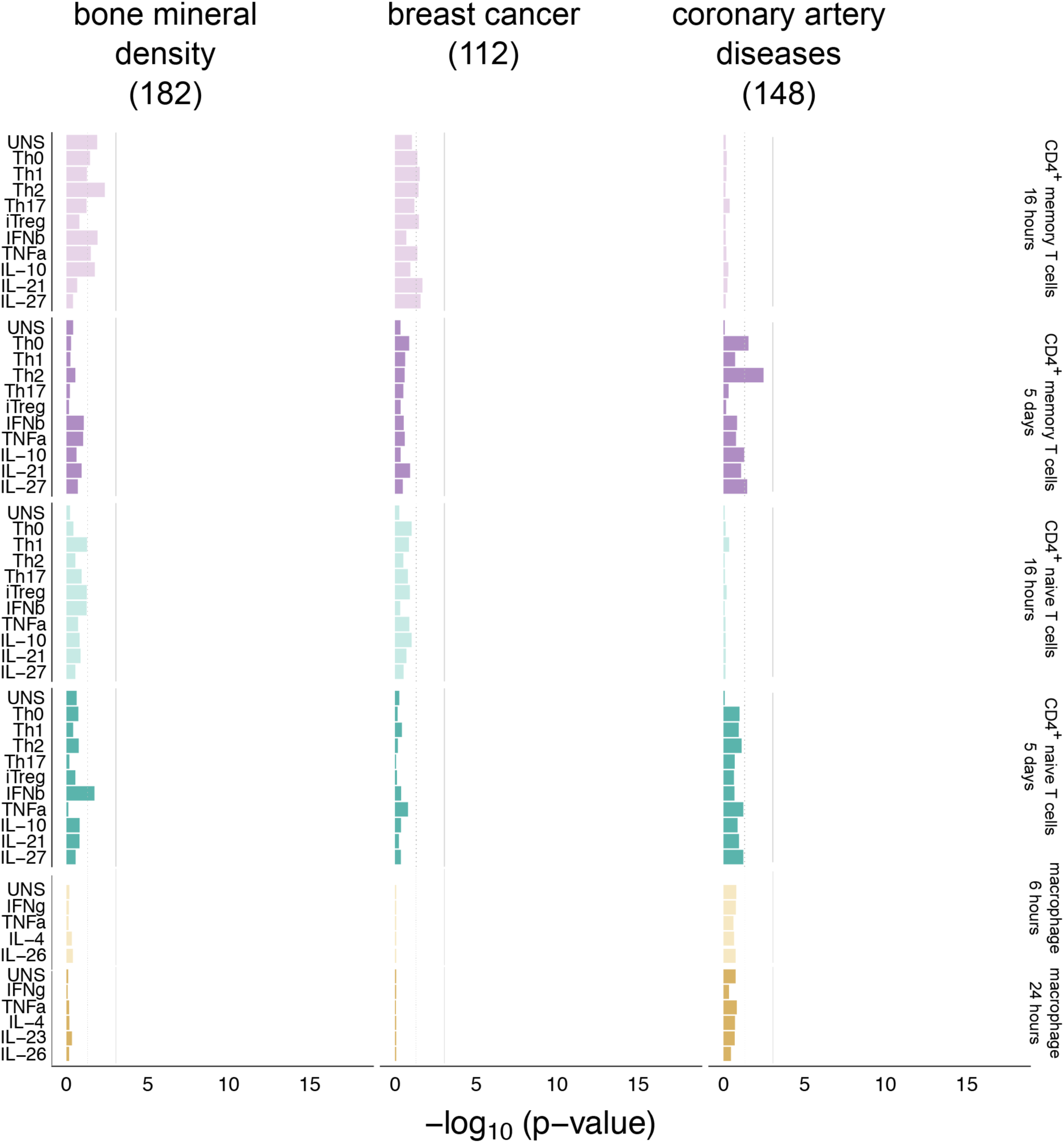
*CHEERS* SNP enrichment in H3K27ac regions in cytokine induced cell states for non-immune traits. The numbers in parentheses represent the number of overlapping peaks per disease. The dotted gray line represents the nominal p-value threshold of 0.05, while the solid gray line represents the Bonferroni corrected p-value threshold of 0.00091 (corrected for the number of cell states in the study).

**Figure S5.**
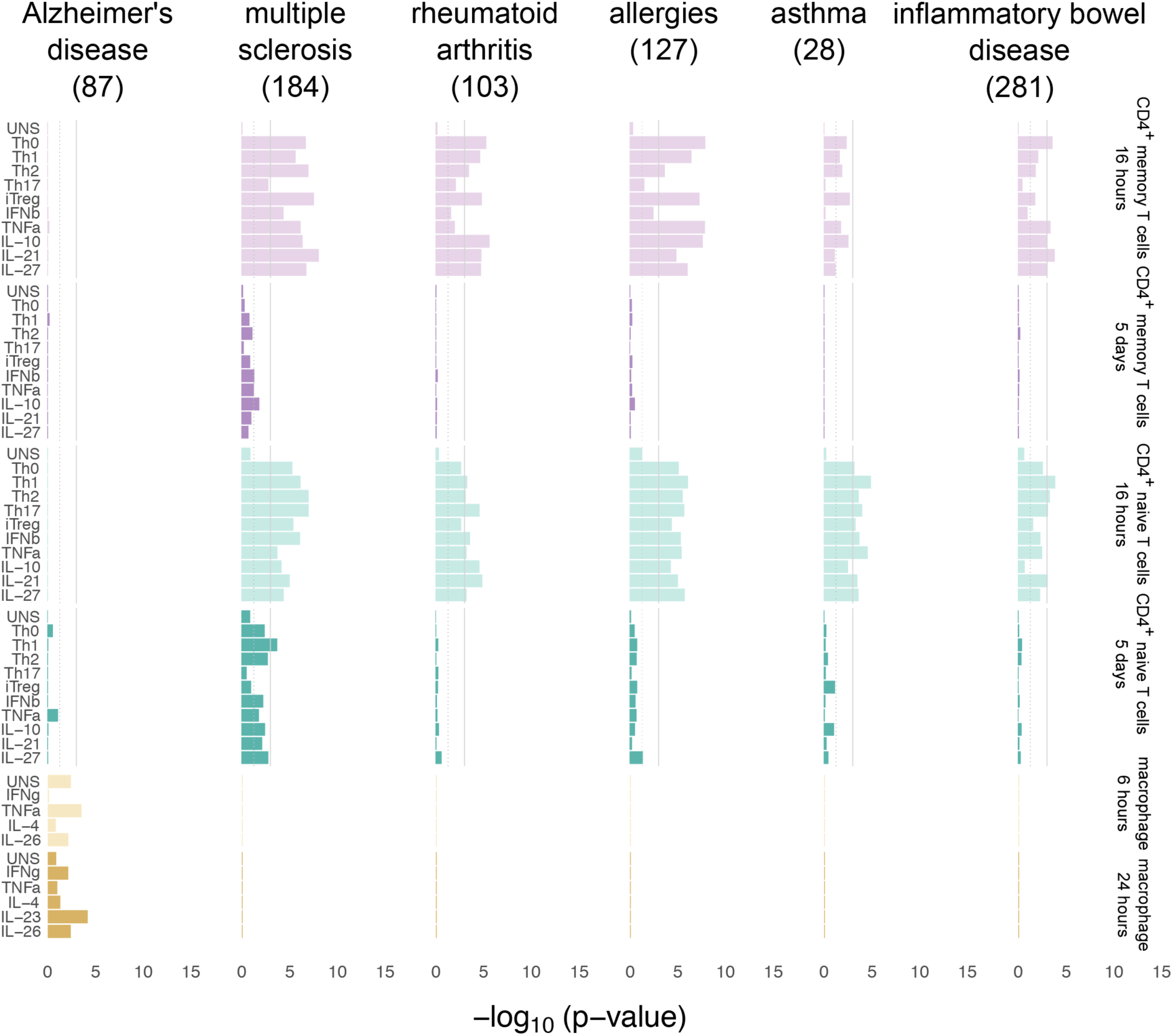
*CHEERS* SNP enrichment in open chromatin regions in cytokine induced cell states. The numbers in parentheses represent number of overlapping peaks per disease. The dotted gray line represents the nominal p-value threshold of 0.05, while the solid gray line represents the Bonferroni corrected p-value threshold of 0.00091 (corrected for the number of cell states in the study).

